# High-throughput single-cell sequencing of multiple invertible promoters reveals a strong determinant of bacterial population heterogeneity

**DOI:** 10.1101/2022.10.31.514637

**Authors:** Freeman Lan, Jason Saba, Yili Qian, Tyler Ross, Robert Landick, Ophelia S Venturelli

## Abstract

Population heterogeneity can promote bacterial fitness in response to unpredictable environmental conditions. Human gut symbiont *Bacteroides* spp., displays variability in single cell surface architectures by combinatorial regulation of promoter inversions that drive expression of capsular polysaccharides (CPS). Using high-throughput single-cell sequencing, we reveal population heterogeneity generated through combinatorial promoter inversion and show that *B. fragilis* populations can access diverse CPS promoter states. Combining our data with stochastic computational modeling, we demonstrate that the rates of promoter inversion regulated by a broadly conserved serine recombinase are a major mechanism shaping population heterogeneity. Exploiting control over the expression of the recombinase, we devise a method for creating phase-locked variants and show that populations with different initial compositions converge to a similar steady-state composition over time. Our approach can be used to interrogate single-cell phase variable states of diverse microbes including bacterial pathogens.

**Summary sentence:** High-throughput single cell sequencing of phase variation reveals regulation as a major driver of population diversification

## MAIN TEXT

Gene regulation in bacteria is shaped by stochastic and deterministic processes. When faced with changing and uncertain conditions, establishing a phenotypically diverse population *a priori* to “hedge” against different environmental stresses can promote population survival. For example, the establishment of persister populations in bacterial pathogens can promote survival under antibiotic treatment (1). The composition (fraction of the population occupying each state) of these heterogeneous populations is influenced by the underlying regulatory mechanisms. Understanding how bacteria regulate the phenotypic heterogeneity at the population-level can reveal important insights into their mechanisms for survival and adaptation.

Phase variation is a ubiquitous mechanism for generating phenotypic heterogeneity in host-associated bacteria such as pathogens and symbionts. (2–8). This process works through diverse mechanisms, such as reversible DNA recombination, to create subpopulations with distinct patterns of gene expression (9). In symbiotic *Bacteroides spp.,* phase variation of capsular polysaccharide (CPS) generates specialized and multi-functional subpopulations having alternative propensities for biofilm development (10), resistance to antibiotics (11), and protection from diverse phages (11,12).

Despite the importance of phase variation to bacterial fitness in diverse environments, limitations in technology have hindered a fundamental understanding of the landscape of single-cell states generated by combinatorial phase variation. Additionally, previous studies of phase variation used fluorescent reporters that disconnect the loci from their natural genetic context and limit the number of loci that can be observed simultaneously (4,13–15). Furthermore, fluorescent reporters produce signals that are temporally disconnected from phase variation due to time delays in fluorescent protein maturation and decay (16).

In the abundant and prevalent genus in the human gut microbiome, *Bacteroides spp.,* a broadly conserved serine recombinase controls promoter inversion at 7–13 CPS biosynthetic loci (16–18). *Bacteroides fragilis (B. fragilis),* a key symbiont, has seven invertible CPS loci (A, B, D, E, F, G, and H), the orientations of which determine the landscape of surface architectures for single cells. An eighth CPS promoter C is constitutively in the ON state to inhibit formation of lower fitness acapsular subpopulations. Omitting C for conciseness in labeling, promoter orientations define discrete ON/OFF states of distinct CPS synthesis operons (e.g., A-E--- denotes A and E in the ON state and B, D, F, G, H in the OFF state) (**Figure 1A**).

**Figure 1.**
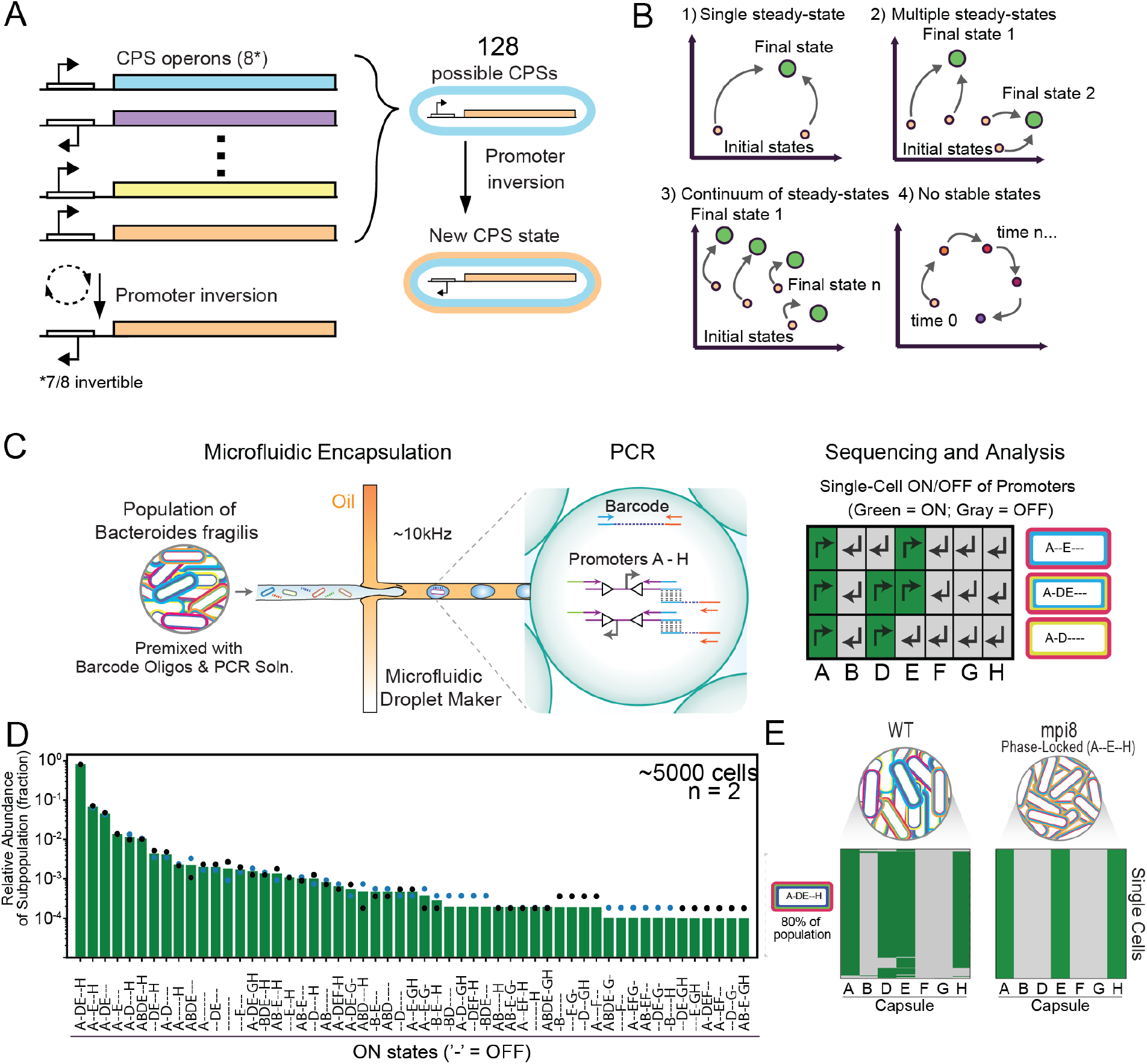
High-throughput single cell sequencing of capsular polysaccharide promoters (CPS) in a key human gut symbiont *Bacteroides fragilis (B. fragilis)* reveals bacterial subpopulations defined by multiple phase variable loci. **(A)** Summary schematic of *B. fragilis* CPS loci. Promoter inversions generates diversity in CPS expression. **(B)** Cartoon schematic of the promoter state landscape of the CPS system under continuous growth. Populations could 1) converge to a single stable state, 2) converge to multiple stable states, 3) exist in a continuum of stable states, or 4) remain in flux with no stable states. **(C)** Cartoon representation of the ultrahigh-throughput single-cell targeted sequencing workflow for *B. fragilis* CPS promoters. We first premix cells from a colony or liquid culture with barcode oligos, high-fidelity PCR mix, and target-loci primers (14 primers for the 7 invertible CPS promoters) containing adaptor sequences that overlap the barcode oligo. Cells in this mix are then encapsulated into picolitre emulsion droplets at limiting dilution so that most droplets contain 1 or 0 cell according to a Poisson distribution. The barcode oligos also exist at limiting dilution so that droplets containing cells also contain 1 or 0 barcodes. Droplets that contain both a barcode oligo and a cell are selectively amplified and linked to genomic loci via overlap extension PCR. The cell is lysed during the initial PCR denaturing step. Only amplicons linking the barcode and loci can be sequenced. After DNA library preparation, sequencing reveals the promoter orientations of all targeted loci for thousands of cells. Right: promoter orientation heatmap for 3 representative cells. Coverage in the ‘on’ orientation for a particular CPS promoter is green. Dashes (‘-’) represent promoters that are oriented off. **(D)** The CPS promoter states of a single colony of *B. fragilis*, shown as a bar plot. Data points represent two technical replicates. **(E)** Heatmaps as described in (A) for thousands of cells, with groups of rows clustered to reflect subpopulations with alternative combinatorial promoter orientations. A control strain with the primary recombinase gene (*mpi*) deleted displays a population with 100% of cells having the expected promoter configuration (A--E--H) for this particular isolate.

Because tools to quantify multiple promoter states within a large number of single cells have not been available, fundamental questions about combinatorial phase variation in any bacterial species remain unanswered. Here, we use a new single-cell sequencing method to investigate the contribution of regulation to combinatorial CPS promoter states within a *B. fragilis* population. Is the population composition of CPS states tightly constrained or loosely influenced by the intrinsic mechanisms of phase variation (in this case, rates of recombination)? In this system, what is the landscape of promoter state compositions that are generated by phase-variation (**Figure 1B**)? Are there one or multiple steady-state compositions or is the system dominated by stochastic processes such that no particular population composition is strongly favored? Can the inversion rates be resolved using single-cell time-series measurements of single-cell states across the population and are these rates coupled or independent? A more detailed and quantitative understanding of this system could provide insights into the contributions of stochastic processes, regulatory network architectures, and the role of history-dependence in shaping the dynamic behaviors of phenotypic diversification by combinatorial phase variation.

### High-throughput single-cell sequencing of multiple invertible promoters

To investigate *B. fragilis* population diversification at the single-cell level, we used an ultrahigh-throughput, single-cell sequencing method referred to as Droplet Targeted Amplification sequencing (DoTA-seq) (19). DoTA-seq can analyze combinations of genes in diverse gram-positive and gram-negative bacteria at the single-cell level. Here we simplified the protocol for profiling the 7 CPS promoter ON/OFF states in *B. fragilis* (gram-negative) **(Methods)**. Briefly, single *B. fragilis* cells and random barcode oligos are encapsulated at limiting dilution into water-in-oil droplets at ~10 kHz rates along with multiplex PCR primer sets (**Figure 1C**). Multiplex PCR amplifies the seven CPS promoters tagged separately with a barcode sequence unique to each droplet to generate a single-cell amplicon sequencing library. The primer-annealing sites flank the inverted repeat sequences where promoter inversion occurs for each operon (17). Thus, each of the seven pairs of primers report the ON/OFF status of their respective promoter through the unique sequence of the amplicon generated (see Methods for detailed breakdown of the quality control and analysis pipeline, and **Supplementary Table 1** for a summary of the number of cells sequenced and passing quality controls for each library).

To determine the technical reproducibility of this method, we sequenced a single colony grown for 48 hours on an agar plate in technical replicates (~5000 cells per replicate). We detected 55 of 128 possible single-cell promoter states within this population, which represent 43% of all possible single-cell promoter states (**Figure 1D**). In a modified strain with the recombinase *(mpi)* responsible for inversion deleted (mpi8) (17), the entire population displayed the same single-cell promoter state (100% of ~9000 cells) (**Figure 1E**). Therefore, the deletion of *mpi* locked the orientation of the promoters across the population. The variation between technical replicates was low for most single-cell promoter states, but increased for states with low relative abundance. Consistent with this result, the coefficient of variation for technical replicates displayed a moderate negative correlation with the number of cells sequenced for each subpopulation **(Supplementary Figure 1)**. This implies that some fraction of the variance could be attributed to random variation due to sampling (i.e. low abundance sub-populations may be inconsistently observed).

### Patterened development of population composition in wild type B. fragilis colonies

After establishing the robustness of the method, we investigated the variation in population-level promoter compositions across multiple colonies (populations derived from single cells) (**Figure 1A**). To this end, we examined single-cell promoter orientations in five wild-type colonies picked at random, representing five independent populations **(Figure 2A)**. Across all five colonies, we observed 77 of 128 possible CPS combinations. Each colony displayed markedly different population compositions with 0–4 invertible promoters oriented in the ON state (**Figure 2B**).

**Figure 2.**
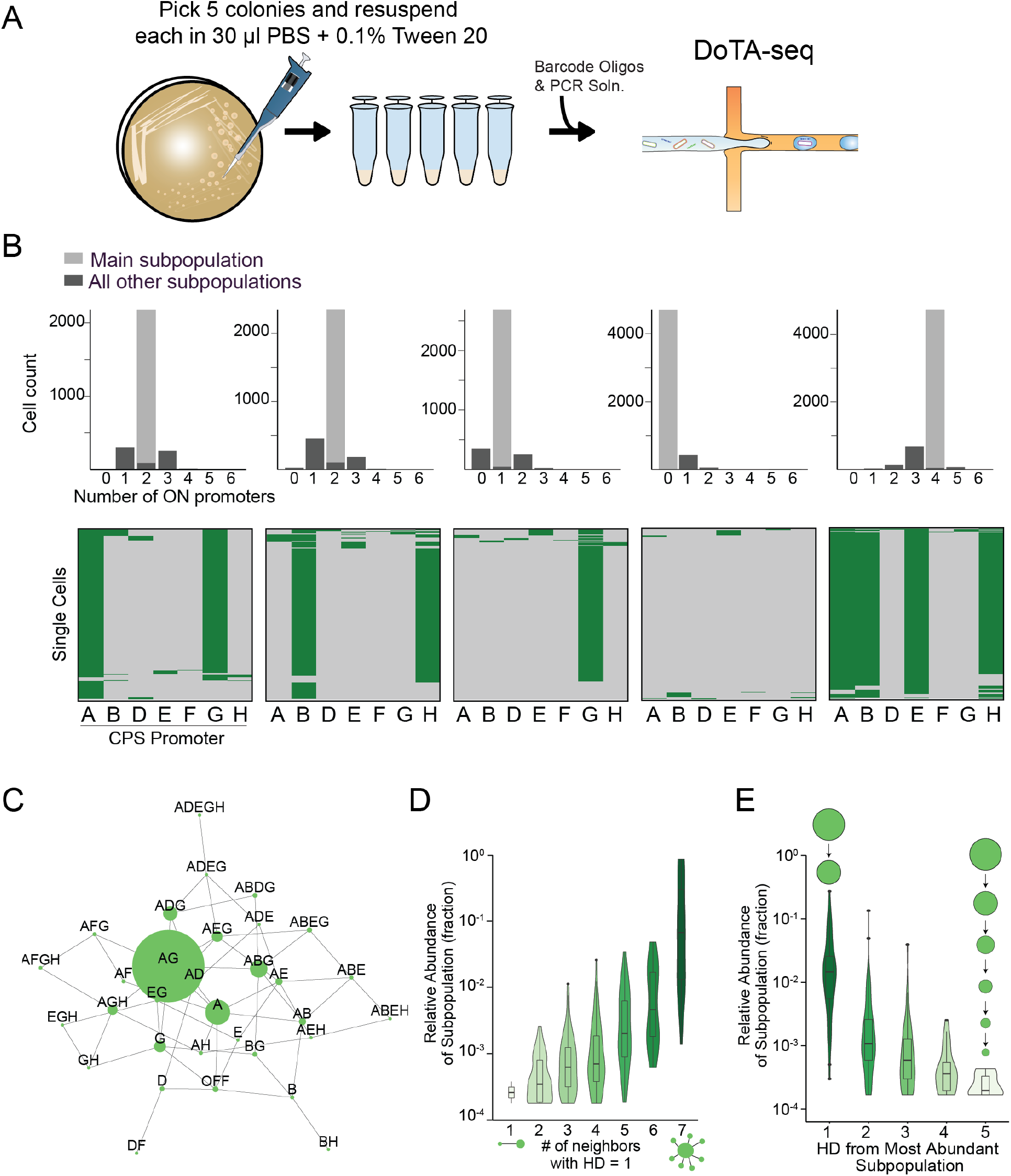
High-throughput single cell sequencing of promoter states in *B. fragilis* colonies reveals diverse compositions with similar features. **(A)** Cartoon representation of experiment. A single glycerol stock was streaked onto an agar plate, and 5 colonies were picked for sequencing after 48 hours of growth. **(B)** Heatmaps of promoter states for 5 wild-type colonies. Top: Histogram of cells having *n* promoters simultaneously oriented on. The light gray bar represents the size of the most abundant state (i.e. main population). **(C)** Representative network representation of single-cell promoter states of colony 1. By connecting subpopulations with a Hamming Distance (HD) of 1 (i.e., separated by a single promoter inversion event), we can generate a undirected network representation of the subpopulations. Each node represents one subpopulation. Node size represents relative abundance. For network graphs of all colonies, see Supplementary Figure 2. **(D)** Violin plot of the distribution of subpopulations with different numbers of neighbors (where HD = 1). All colonies show the characteristic main subpopulation surrounded by progressively lower abundance subpopulations. **(E)** Violin plot of the distribution of subpopulations versus the distance from the most abundant subpopulation showing that most of the large populations are found close (lower HD) to the main subpopulation.

To understand the global properties of the population compositions across each colony, we generated an undirected network that represents the relationships between single-cell promoter states (**Figure 2C** and **Supplementary Figure 2**). Each single-cell promoter state is represented by a node proportional in size to its relative abundance in the population. Nodes are connected if their Hamming distance (HD) is equal to one (i.e. separated by a single promoter-inversion). For example, the HD between combinatorial states A--E--- and A-DE--- is 1.

Aggregate analysis of the networks for all colonies revealed patterns in population composition. While subpopulations with many (5–7) promoters turned off were frequently observed, subpopulations with many (5–7) promoters simultaneously on were rare. This pattern suggests that *B. fragilis* limits concomitant promoter-ON orientations. In addition, all observed subpopulations (accounting for ~22-34% of possible single-cell states) are connected (1 HD away) to at least one other observed subpopulation. This is consistent with the possibitility that the rates of inversion are much slower than the rate of cell division. Further, larger (i.e., higher abundance) subpopulations had many (>6) immediate (HD = 1) neighbors **(Figure 2D)**. For comparison, most other lower abundance subpopulations were connected to ~3–5 immediate neighbors **(Supplementary Figure 3A)**. This pattern is consistent with the notion that high abundance populations arose earlier in population development and gave rise to their lower abundance neighbors.

Higher abundance subpopulations were closer (i.e., lower HD) to the most abundant subpopulation **(Figure 2E**, Spearman’s Rho −0.65, p < 2.2e-16 between subpopulation size and HD). Finally, most of the cells were only one promoter inversion away from the most abundant promoter state (main promoter state) **(Figure 2B,E)**. These observed patterns in population-level promoter state network properties could arise from two possibilities. First, the population composition reflects a snapshot in early stages of a diversifying population, where the major subpopulations are more closely related to the single-cell promoter state of the initial founding cell. Alternatively, the observed promoter-state composition reflects the longer-term steady-state to which each colony converged to independently.

### Populations with different initial promoter state compositions display converging trajectories over time

To determine whether these populations are in early stages of diversification or reflect multiple steady-state compositions, we individually cultured the five colonies for two weeks in liquid media (**Figure 2B**). The cultures were diluted every 8 or 16 hours alternately with different inoculum volumes to prevent the cultures from entering late stationary phase (**Methods**). To track the temporal diversification process, we performed DoTA-seq on the populations on day 3, 7, and 14 **(Figure 3A)**. The Shannon diversity of individual cultures increased over time as new and distinct single-cell promoter states arose, and the population became more evenly distributed among different combinatorial promoter states **(Figure 3B)**. By day 14, each population exhibited on average 89 +/- 9 unique single-cell states and displayed 117 unique single-cell states across all colonies **(Supplemental Table 2)**. Although these populations had not yet reached steady-state (i.e. population-level promoter composition was still changing over time), the population compositions became increasingly similar to each other as time progressed (**Figure 3C**). These results suggest that populations from different initial starting states eventually converge to a single steady-state population composition. However, the time for convergence to a steady-state was on the scale of weeks, which suggests that weak forces shape the population composition towards a single preferred state under our culture conditions. In addition, the large number of single-cell promoter states detected in the populations on day 14 highlights the diverse repertoire of variation available to *B. fragilis* populations to cope with environmental stresses.

**Figure 3.**
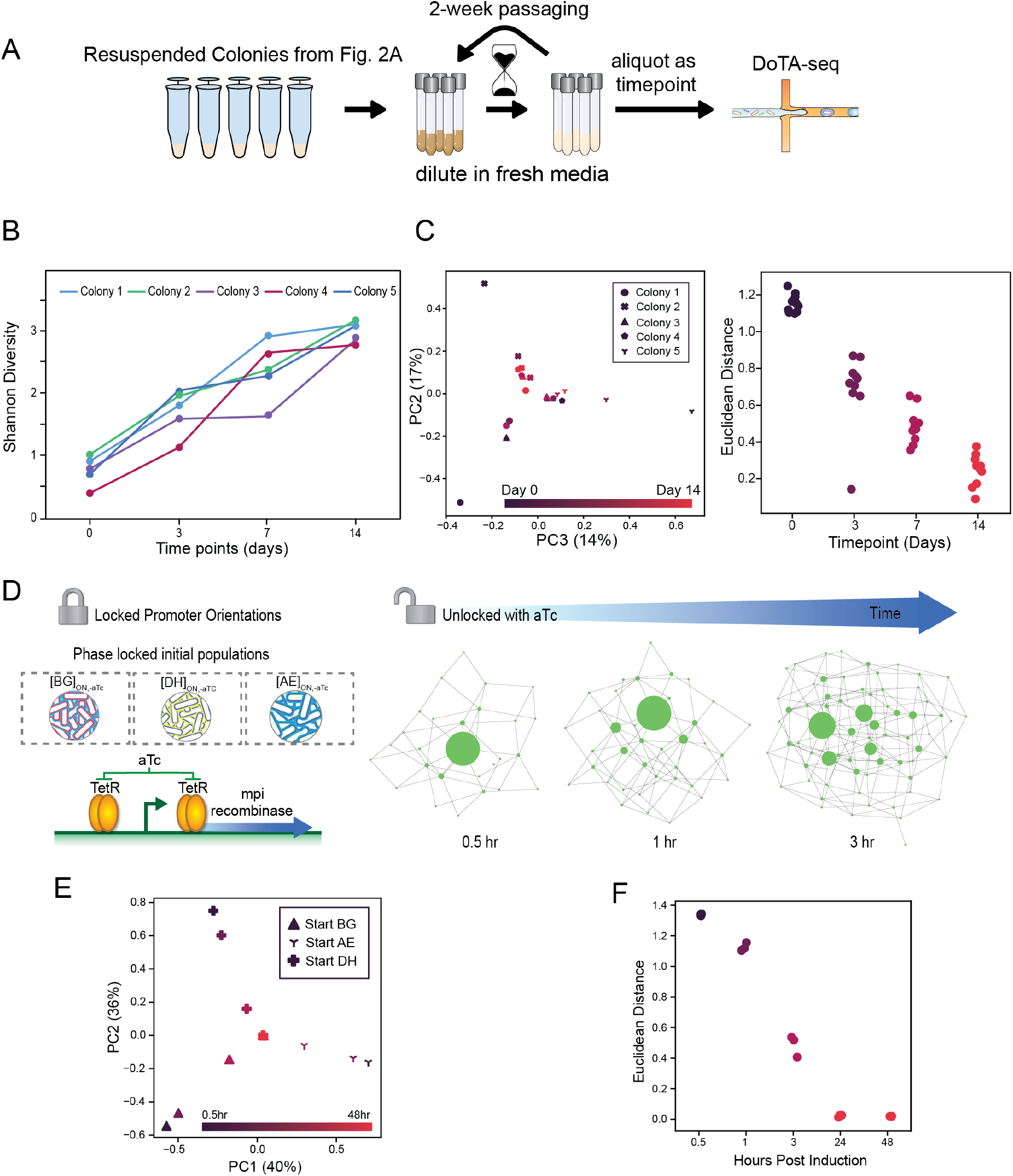
Population-level promoter state composition converges to a single steady-state composition over time. **(A)** Cartoon schematic of the experiment. The 5 colonies characterized in Figure 2 were grown in liquid media for 14 days and diluted periodically to prevent culture from reaching late stationary phase. Samples are taken and promoter states were characterized using high-throughput single-cell sequencing at multiple timepoints. **(B)** Shannon diversities of the 5 wildtype populations originating from the 5 colonies over each timepoint. Shannon diversity increases with time for all colonies. Day 0 represents the populations as colonies on a plate. **(C)** The populations show a converging trajectory over time to a single steady-state. Right: Population-level promoter state compositions are visualized bty principal component analysis (PCA). Marker shapes represent different populations. Marker colors represent different timepoints. Left: The pairwise Euclidean distance between all 5 populations at each timepoint. The trend of decreasing Euclidean distance shows a converging trajectory of the populations to a single steady-state. **(D)** Cartoon representation of the induced promoter inversion experiment. The native recombinase gene (*mpi*) responsible for promoter inversions was driven by an aTc-inducible promoter. Inversion is induced in phase-locked variants during log phase by the addition of 100 ng/μL aTc into the cultures. **(E)** aTc-induced cultures also show a converging trajectory based on PCA. Marker shapes represent different cultures. Marker colors represent different timepoints. **(F)** Pairwise Euclidean distances between the 3 aTc-induced populations at different timepoints post induction. By 24 hours, the 3 populations converged to a steady-state.

The population dynamics and the steady-state promoter distributions could be driven by regulation (e.g., promoter inversion rates), variation in subpopulation fitness, or a combination of these factors. We first considered the contribution of the recombinase-mediated promoter inversion rates to the population dynamics. In many bacterial species, the rates of phase-variation can change in response to environmental cues (4). In *B. fragilis* specifically, expression of the Mpi recombinase may be regulated by additional phase-variation mechanisms (20). To eliminate these potential additional layers of regulation, we constructed a strain enabling synthetic control of expression of the recombinase. To this end, we introduced a tightly regulated (i.e., low leaky expression), aTc-inducible *mpi* at an ectopic location on the genome in an *mpi* deletion background in *B. fragilis*. In the presence of high aTc concentrations, *mpi* can be expressed at high levels from this inducible promoter **(Fig. 3D)**.

We first used this system to generate and isolate phase-locked variants (**Supplementary Figure 5**) in different single-cell promoter states (A--E---, -B---G-, and --D---H) in the absence of aTc. In the presence of aTc, recombinase expression is turned on and the promoter states diversify across the population as a function of time. To investigate the diversification process over time, we cultured the phase-locked strains to mid-exponential-phase, induced recombinase expression, and then sampled these populations over time after induction (**Figure 3D**). The induced populations showed converging population composition trajectories (**Figure 3E**) and approached a single steady-state population composition 24 hours following induction of recombinase (**Figure 3F**). Thus, high expression of recombinase resulted in a fast convergence of the population to the steady-state composition, imposing a strong constraint on population composition.

To evaluate the contribution of fitness differences for each variant to the observed population dynamics, we determined the growth rates of the phase-locked variants under our culturing conditions. We did not observe statistically significant differences in growth rates between different CPS locked strains when grown in our culturing medium (p > 0.19, one-factor ANOVA) (**Supplementary Figure 4)**. This suggests that variation in subpopulation fitness was not a strong contributor to the population dynamics in our experimental conditions.

### Computational modeling reveals a major mechanism governing population dynamics

Computational modeling is a powerful way to gain mechanistic insight into complex biological systems (21). However, the development of computational models can be challenged by the complexity of biological systems combined with a lack of high quality data needed to sufficiently constrain model parameters. Single cell data can provide constrain stochastic model parameters better than bulk measurements (22). Leveraging our single-cell data, we constructed a stochastic mechanistic model to describe the population composition as a dynamic process in which promoters flip independently and stochastically at the single-cell level. This process is represented as a continuous-time Chemical Master Equation (CME) model consisting of 128 discrete states (**Figure 4A, Supplementary Materials**). The parameters of this model were estimated by fitting the analytical solution of the CME to the time-series single cell data (**Methods**).

**Figure 4.**
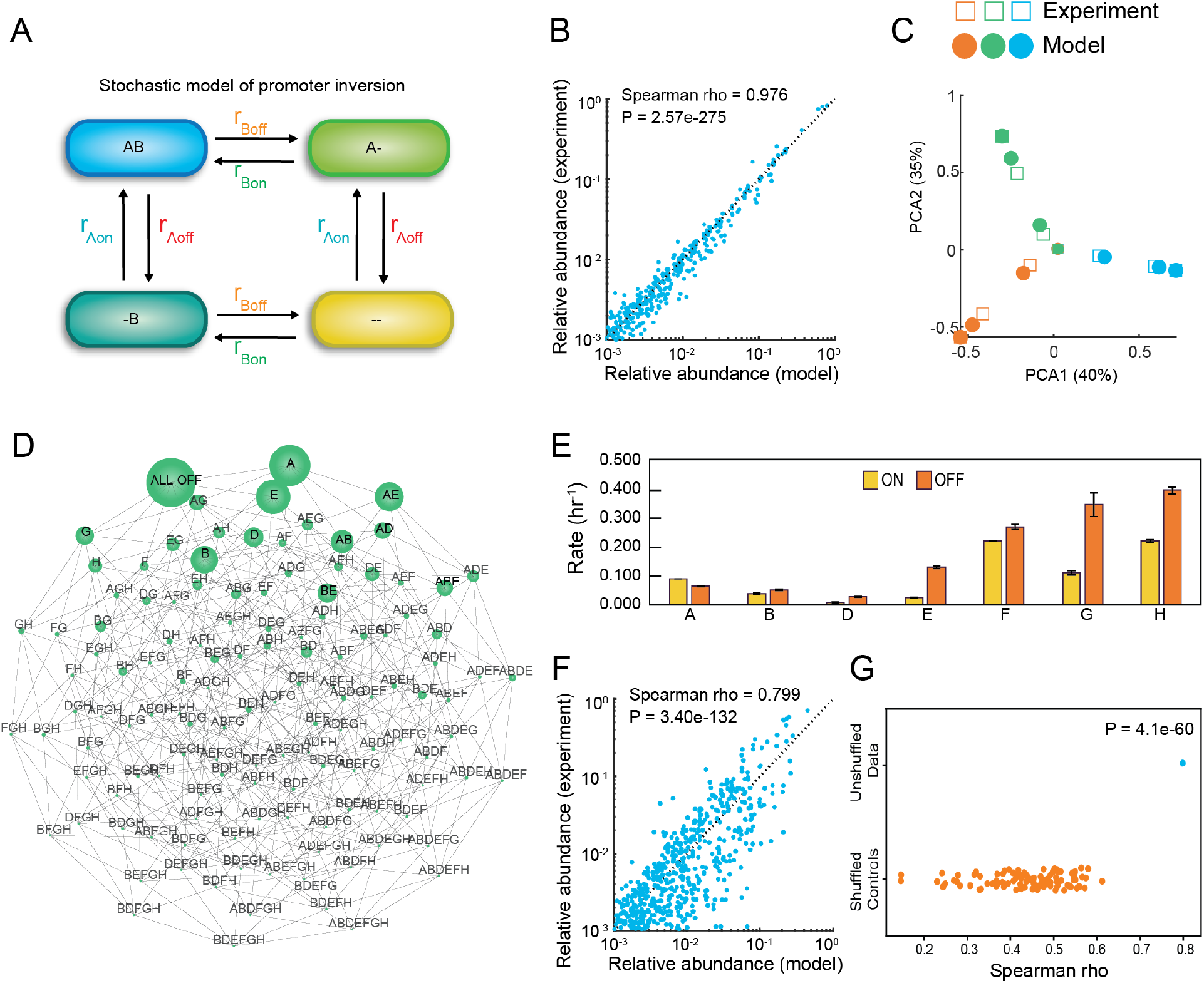
A stochastic dynamic model of promoter inversions demonstrates that promoter inversion rates are a major variable shaping the temporal changes in promoter state diversification. **(A)** Schematic (only two promoters) of the stochastic dynamic computational model for capturing temporal changes in single-cell promoter states. In the simplified representation, the cell transitions between four different states by inverting promoters on and off with different kinetic rate parameters. A and B represent different operon promoters in the ON state and “-” represents the promoter in the OFF state. In the model, there are 7 promoters, 14 kinetic parameters, and 128 different states. **(B)** Scatter plot of promoter state relative abundance of the 128 possible promoter combination states predicted by model versus experimentally measured for each time point of the aTc-inducible experiment. Spearman rho represents the Spearman correlation coefficient. **(C)** Model predictions and experimental data visualized using Principal Component Analysis (PCA). **(D)** Undirected network graph of the model’s prediction of the promoter state composition. Subpopulations are connected by an edge if their hamming distance is equal to one. Node size represents relative relative abundance of the subpopulation. **(E)** Inferred kinetic rate parameters for promoter inversions derived from the model fitted to the aTc-inducible time-series data. Error bars represent 1 s.d. of the Monte-Carlo Marko chain parameter distribution. Posterior parameter distributions are shown in Supplementary Figure 7. **(F)** Scatter plot of promoter state relative abundance of the 128 possible promoter combination states predicted by model versus experimentally measured for each time point for the 14-day wild-type experiment. **(G)** Categorial scatter plot of the Spearman’s correlation of a model fit to the label shuffled controls versus wild-type time-series data.

We fit the model to the single-cell time series data to evaluate the contribution of promoter inversion rates to the observed population dynamics. This model fit the time series measurements of promoter states in the *mpi* synthetically regulated strain extremely well (Spearman’s Rho ~0.98, p = 2.57e-275), suggesting that the assumptions of the model were consistent with the experimental observations (**Figure 4B-C**). In addition, all model parameters were well-constrained by the time series data except the promoter inversion rates D-ON and G-OFF. The posterior parameter distributions for these parameters displayed coefficients of variation greater than 5%, indicating that these parameters were moderately constrained **(Supplementary Table 3)**. To determine if the observed goodness of fit was due to the patterns observed in the experimental data, we randomly shuffled the dataset labels and re-fit the CME model. The model did not fit well to the label-shuffled datasets **(Supplementary Figure 7)**. In sum, the population diversification trajectories of the *mpi* synthetically regulated strain are driven primarily by independent and differential promoter inversion rates.

To investigate the longer-term behavior of the population beyond the experimentally measured time points, we determined the steady-state of the CME model (**Figure 4D**). The steady-state exhibits high diversity, with most single cells displaying 0-2 invertible promoters simultaneously ON. In addition, the steady-state is dominated by promoter combinations with promoter A in the ON state. Dominance of the promoter A in the ON state is reflected in the inferred promoter inversion rates, where only promoter A has a higher rate of flipping ON than OFF (**Fig. 4E)**. Promoters F, G, and H display higher inversion rates (both ON and OFF) than the other promoters, leading to a faster convergence of these promoters to steady state than the other promoters with smaller magnitude ON and OFF rates **(Supplementary Figure 8)**.

The observed differences in kinetic rates could reflect different roles that CPSs play in the biology of the bacterium. For example, capsular polysaccharide A (PSA) has been shown to be essential for colonization (23), pathogenesis (24), and was found on average to be oriented ON in more than 50% of WT populations in every tested condition (17,25). This is consistent with the inferred higher ON than OFF rate for promoter A. Capsular polysaccharide G (PSG) was previously shown to be highly context dependent, only detected in high proportions in the context of a microbial community *in vivo* (25). Here, promoter G had faster kinetic rates but was biased towards the OFF state. Thus, the ON and OFF rates may influence sensitivity of *B. fragilis* CPS compositions to environmental conditions.

To evaluate the contribution of promoter inversion rates to the population dynamics of wild-type *B. fragilis,* we fit the CME model to time-series measurements of the 5 wild-type populations **(Figure 3A)**. The model displayed a reasonable fit to these data (Spearman’s Rho ~ 0.8, p = 3.40e-132) **(Figure 4F)** and the fit was substantially better than the label-shuffled control data (p = 4.1e-60, one sample t-test) **(Figure 4G)**. This result implies that the independent rates of promoter inversion could explain a portion of the dynamics in this experimental system. The posterior distributions for certain parameters (F-OFF and G-OFF) exhibited bimodality and were not well-constrained (>5% coefficient of variation in parameter distribution). This suggests that additional factors beyond those captured in the model (such as regulation of recombinase levels or activity) contribute to the population dynamics in wildtype populations, especially for promoters F and G (**Supplementary Figure 6B, Supplementary Table 3**). Consistent with this notion, promoters F and G harbor extended inverted repeat DNA elements surrounding the promoter (26), potentially influencing recombinase activity or dependence on other regulatory factors. Although Mpi is required for inversion, Mpi may interact with other regulatory proteins to modulate inversion at different loci, providing an additional layer of regulation over specific CPSs (23). Further work is required to elucidate these additional layers of regulation of promoter inversion rates, especially in physiologically relevant environmental contexts (20).

In sum, the model’s excellent fit to time series measurements of the *mpi* synthetically regulated strain indicates that independent promoter inversion rates are a major mechanism driving the population-level CPS promoter state composition of *B. fragilis* populations. By contrast, the model’s moderately good fit to the wildtype data suggests that additional layers of regulation are present, especially for promoters F and G.

### Summary and outlook

Phase variation has been shown to be a wide-spread mechanism of generating population heterogeneity for diverse bacteria (3,4). However, we have a limited quantitative understanding of the diversification process, the landscape of single-cell states and the driving regulatory networks. By profiling the phase variable CPS loci in *B. fragilis* populations at the single-cell level over time, we uncovered the quantitative contribution of promoter inversion rates to the observed dynamic behaviors. The population dynamics are fundamentally shaped by different rates of promoter inversion, which influences the steady-state population composition and how quickly the population reaches the steady-state. The convergence of this system to a single-steady-state is in contrast to other diversification gene regulatory networks, which are strongly influenced by interlinked positive and negative feedback loops and exhibit bistable behavior (27). These results provide a foundation for further understanding a human gut symbiont in the context of their natural life cycles, such as through colonization of the human gut. This novel method of studying phase-variation, as well as our findings can be extended to other diverse bacteria, including pathogens, that use phase variation in their life cycle (2,28–35). Thus, our results set the stage for studying how population heterogeneity is used by bacteria to respond to environmental perturbations such antibiotic stress, microbial warfare, and phage and host immune cell attack.

## METHODS

### Plasmids and Strains

*B. fragilis* NCTC9343 was obtained from ATCC. Lyophilized culture was resuspended in SBM and frozen in 25% glycerol. All *B. fragilis* cultures were grown at 37°C in an anaerobic chamber (Coylabs) with an atmosphere of 2.5 ±0.5% H2, 15 ±1% CO2 and balance N2. We note that production of a stock following outgrowth single colony, rather than directly from lyophilized culture as done here, will generate a stock with significantly lower initial diversity.

To generate the inducible recombinase strains locked under different promoter orientations, we integrated chromosomally the *mpi* recombinase gene under the control of a tetracycline inducible promoter in a *mpi* deletion strain. pNBU2_erm-TetR-P1T_DP-GH023 was a gift from Andrew Goodman used for creating the tetracyline-inducible recombinase strain (Addgene plasmid # 90324; http://n2t.net/addgene:90324; RRID:Addgene_90324). Briefly, we cloned mpi (also known as ssr2) into pNBU2_erm-TetR-P1T_DP-GH023 via Gibson assembly, transformed into E. coli strain RL1752 via electroporation, and sequence-verified using whole-plasmid sequencing (pJS041). pJS041 was next transformed via electroporation into the donor strain E. coli BW29427 and plated on LB agar containing 100 μg/mL ampicillin and 150 μM DAP. Overnight cultures were created by inoculation of a single colony of donor (pJS041/E. coli BW29427) or recipient (Δmpi M44) colonies into LB containing 100 μg/mL ampicillin and 150 μM DAP or Supplemented Basal Medium (SBM), respectively. In the morning, 10 μL of overnight donor culture was used to inoculate 5 mL LB containing the same supplements, while 250 μL of overnight recipient culture was used to inoculate 20 mL of SBM (2% proteose peptone, 0.5% yeast extract, 0.5% NaCl, 1.5% agar, supplementing 0.5% glucose, 0.5% K2HPO4, 0.05% cysteine, 5 μg/mL hemin, and 0.25 μL/100 mL vitamin K1 after autoclaving). The donor culture was grown to mid-late log phase while recipient culture was grown mid-log phase, then the donor culture was pelleted by centrifugation at 4000xg for 5 minutes, washed by resuspension in 5 mL LB and centrifugation. The donor pellet was then resuspended with the 20 mL recipient culture, pelleted at 3000g for 10 minutes, then resuspended in 1 mL standard SOC medium containing 5 μg/mL hemin and 5.5 μM vitamin K1 (SOC+HK), spotted on BHIS (37g/L with 5 μg/mL hemin and 0.25 μL/100 mL vitamin K1) agar containing 150 μM DAP, and incubated upright, overnight, and aerobically at 37°C. The next day, matings were scraped from plates and resuspended in 2 mL of SOC+HK. Dilutions were plated on BHIS agar containing 200 μg/mL gentamicin and 5 μg/mL erythromycin. Colonies were screened for their ability to express *mpi* under induction of 100 ng/μL anhydrotetracycline by SDS-PAGE.

### Plating and harvesting of B. fragilis colonies

Cells were streaked out directly from frozen glycerol stock at the lab bench on to SBM agar plates and the plates were incubated for two days (>36 hours) before colonies were harvested using a sterile pipette tip and resuspended in PBS + 0.1% Tween 20 by pipetting up and down.

### Long-term culturing of *B. fragilis* populations

Five colonies were resuspended in 30 μL PBS + 0.001% Tween20. 10 μL of each suspension was used to inoculate separate 1 mL overnight cultures. The remaining suspension was used for microfluidic analysis of colonies. In the morning, timepoints were taken from each of the five cultures: each culture was briefly vortexed, then two 100 μL aliquots were mixed with equal volumes of 50% glycerol (technical replicates) and flash-frozen in a dry ice-ethanol bath. Each day for 2 weeks, cultures were passaged twice: in the morning (16 hours after inoculation), 10 μL of the overnight culture was used to inoculate 1 mL of SBM (1:100); in the evening (8-9 hours later), 1 μL of saturated culture was used to inoculate 1 mL of SBM (1:1000).

### Growth curves of wildtype and CPS promoter locked strains

Overnight cultures of each strain are inoculated from frozen glycerol stocks into SBM media. The next day, the OD600 of each overnight culture is measured on the Nanodrop One (Thermofisher) using the cuvette setting, and a 96-well flat bottom assay plate (Midsci 781602) is prepared with 200 μL of SBM media and each culture is added to two wells to an initial OD600 of 0.01. The plate is incubated in a Tecan F-200 plate reader at 37°C in the anaerobic chamber, taking an OD600 reading every 30 minutes for 48 hours.

### Fitting of growth curves to data

The raw OD600 readings are truncated to remove the death phase and the values normalized to between 0 and 1 for each individual curve. This is to account for potential differences in OD600 per cell for each CPS variant. A logistics growth ODE model (d[X]/dt = (r + alpha-X)*X) where X is the normalized abundance, r is the growth rate, and alpha is the self-inhibitory parameter) is fit to this data using least squares fit function in python. The growth rate is taken as the exponential growth parameter in the logistics model. Scripts used in this analysis can be found on github (See code availability).

### Fabrication of microfluidic droplet maker

Microfluidic masters were fabricated in a clean room using soft lithography (36). SU-8 3000 photoresist (MicroChem) was spun on a silicon wafer (University Wafers) to achieve a thickness of 30 μm. Photolithography masks **(supplementary materials)** were ordered from CAD/ART and used to pattern the photoresist using a UV Led (ThorLabs) for 1 min 45s. The patterned photoresist was baked for 5 minutes at 95oC then developed using SU-8 developer (PGMEA) for 2 minutes then baked at 200oC for 2 minutes.

PDMS devices were created by curing PDMS elastomer (Sylgard-184) at a 1:11 crosslinker to elastomer ratio over the silicon master. These devices were cut out using a scalpel and holes punched using a 0.75mm reusable biopsy punch (World Precision Instruments). PDMS devices were bonded to glass slides using a plasma cleaner (Harris Plasma), and then treated with Aquapel (Aquapel Glass Treatment) to render them hydrophobic.

### Single cell sequencing of *B. fragilis* populations

*B. fragilis* cells were pelleted by centrifugation, washed with 1 mL of PBS + 0.1% Tween 20, then resuspended in 100-500 μL of PBS + 0.1% Tween 20. OD600 readings were taken of the resuspension using the pedestal mode of the Nanodrop One (Thermo Scientific), which is used as proxy for cell concentrations for input into the DoTA-seq workflow.

A PCR reaction mix is prepared as follows: 25 μL Q5 Ultra II PCR mastermix, 1μL of primers mixture consisting of P7(10 μM)-Barrev(1 μM), 0.7μL of Inverton targeting primers mixture (20 μM total), 0.5 μL of Barcode Oligos (1 pM) which has been freshly diluted from 500 pM with TE buffer. See supplementary table 4 for a list of oligonucleotides used in this paper.

This PCR mixture is combined with 25μL of cells diluted with pre-injection buffer (10mM HEPES pH 7.5, NaCl 25 mM, EDTA 0.1 mM, Tween-20 2% v/v) such that the final OD of cells in the final mixture is 0.0025. This mixture is then injected into the droplet making microfluidics device using syringe pumps (New Era) and 1mL syringes (BD) at a flowrate of 400 μL/hr and 800 μL/hr of Biorad Evagreen droplet making oil (Biorad) to generate droplets.

Droplets are collected into a PCR tube (Fisher Scientific #14222292) and thermo-cycled in a Biorad CX-100 thermocycler as follows: 98oC for 2 minutes, and 40 cycles of 98oC and 30 seconds at 65oC for 5 minutes, followed by 72oC for 10 minutes and hold at 12oC.

After PCR, the coalesced droplets were removed using a pipette, and the emulsion was broken on ice by adding 20 μL 500mM EDTA and 20 μL perfluoro-octanol (Sigma 370533), then vortexed followed by a spin pulse centrifugation. The aqueous phase was transferred to another tube by pipette and cleaned up using a Zymo cleanup and concentrator kit (Zymo Research D4013), then subject to size selection using SPRI-select beads (Beckman Coulter B23317) using 0.7x volume of beads. A further round of size selection was performed in 100 mM NaOH, 10% Ethanol, and 1.4x volume of beads to increase the purity of the library.

All libraries were sequenced on the Miseq using V3 150cycle kits with custom sequencing primers **(supplementary table 4)**, using 155 cycles for read 1, 18 cycles for the I7 index, and 8 cycles for the I5 index.

### Analysis of sequencing data

The raw sequencing reads are obtained from the Miseq. Read 1 represents the targeted amplicons. The first index read represents the unique cell barcode. The second index read is used to multiplex libraries from different experiments. Demultiplexing of different libraries (index 2) is performed by the Miseq software. Cell barcode demultiplexing and quality control is performed using a custom python script (R4-parser.py). The filtered reads were mapped to custom built reference databases containing B. fragilis CPS promoter sequences and is available on GitHub (see code availability) using Bowtie2 V2.3.4.1(37) using “--very-sensitive” presets. The mapped reads were analyzed using custom scripts as follows: The mapped reads are organized into read groups consisting of reads with the same unique cell barcode representing amplicons from the same droplet. Read groups with too few reads are removed. The filtered groups are transformed into a table containing the barcode and a tally of the number of reads that are mapped to each target (SAM-analysis.py). We further filtered the reads by removing the read groups if the reads mapping to any promoter (sum of ON and OFF orientation) are less than 1% of total reads for that group. Subsequently, if reads in any promoter orientation were less than 1% of total reads for the barcode, we set the value to 0; this accounted for expected noise. We next discretized the data by replacing values with 1 or 0, corresponding to ON or OFF, respectively. We removed barcodes if neither orientation was found for any promoter (i.e., if ON + OFF values = 0). We also removed read groups if both orientations were found for any locus (ON + OFF = 2). The set of remaining read groups are used to determine the frequencies of each promoter state combination in the population **(CPS-analysis.R)**. This processed list of frequencies for each promoter state is available in Supplementary Table 2. All scripts are available on GitHub (see code availability).

### Computational modelling of promoter inversion dynamics

For a single cell, the promoter flipping process is represented mathematically as a continuous-time Markov process consisting of 128 discrete states. In an infinitesimal time interval, the propensity for a cell to switch from state *x* to state *y* with a single promoter change is a product of (1) the flipping rate constant of that promoter and (2) the probability that the cell is currently in state *x*. We assume that this stochastic flipping process is ergodic: the frequency of a subpopulation at time *t* represents the probability of an individual cell to be in this state. This assumption allows us to fit the analytical solution of this model to experimental subpopulation frequencies to infer the flipping rate constants. Parameter inference is performed using a Markov Chain Monte Carlo approach, which accounts for uncertainties in measurements of subpopulation frequencies (ie. subpopulations close to stochastic limit of detection). For a detailed description of the modelling workflow, please see **supplementary materials BF-modelling.pdf**. All scripts are available on GitHub.

## Code availability

All scripts and code used in analysis are available on Github at **https://github.com/lanfreem/Bfragilis-CPS.git**.

## Data availability

Raw sequencing data and OD600 plate-reader data are available on Zenodo at the time of publication.

## Acknowledgements

We would like to thank Professor Laurie Comstock at the University of Chicago for supplying published strains and offering feedback on the preliminary data. J.S. would like to thank Dr. Mike Wolfe for discussion and guidance for data processing. J.S. was supported in part by the NIH Biotechnology Training Grant (T32 GM135066 and T32 GM008349) and NIH F31 Graduate Fellowship (F31GM142153). J.S. is supported additionally by the SciMed Graduate Research Scholars Fellowship – support for this fellowship is provided by the Graduate School, part of the Office of Vice Chancellor for Research and Graduate Education at the University of Wisconsin-Madison, with funding from the Wisconsin Alumni Research Foundation and the UW-Madison. F.L. was supported in part by the Burroughs Wellcome Fund Careers Award in the Scientific Interface. This work was supported by the National Institutes of Health R35 GM124774, R35 GM12477409 and R21 AI156438, and Army Research Office W911NF-19-1-0269.

## Author Contributions

O.S.V, R.L., F.L. and J.S. conceived of the study. F.L. and J.S. conceived and performed experiments and conceived and wrote custom scripts for data analysis and processing. F.L. conceived of and executed the microfluidics technology protocol, and generated sequencing data and sequencing analysis. J.S. conceived of and executed the method for generating and isolating phase-locked variants. F.L., J.S., and T.R. performed microfluidics experiments. Y.Q. conceived and executed computational modeling, wrote custom scripts related to these models and performed analyses of these data. F.L. conceived of and executed other computational models. O.S.V., R.L., F.L., J.S. and Y.Q. analyzed the data. O.S.V., F.L., and J.S. wrote the manuscript. O.S.V., R.L., F.L., J.S. and Y.Q. contributed to the design of Figures. O.S.V. and R.L. supervised the study and secured funding.

## References

1. Stapels DAC, Hill PWS, Westermann AJ, Fisher RA, Thurston TL, Saliba AE, et al. Salmonella persisters undermine host immune defenses during antibiotic treatment. Science. 2018 Dec 7;362(6419):1156–60.

2. Gunther IV NW, Snyder JA, Lockatell V, Blomfield I, Johnson DE, Mobley HLT. Assessment of Virulence of Uropathogenic *Escherichia coli* Type 1 Fimbrial Mutants in Which the Invertible Element Is Phase-Locked On or Off. Infect Immun. 2002 Jul;70(7):3344–54.

3. van der Woude MW, Bäumler AJ. Phase and Antigenic Variation in Bacteria. Clin Microbiol Rev. 2004 Jul;17(3):581–611.

4. Bayliss CD. Determinants of phase variation rate and the fitness implications of differing rates for bacterial pathogens and commensals. FEMS Microbiol Rev. 2009 May;33(3):504–20.

5. Cota I, Blanc-Potard AB, Casadesús J. STM2209-STM2208 (opvAB): A Phase Variation Locus of Salmonella enterica Involved in Control of O-Antigen Chain Length. Chakravortty D, editor. PLoS ONE. 2012 May 11;7(5):e36863.

6. Foley J. Mini-review: Strategies for Variation and Evolution of Bacterial Antigens. Comput Struct Biotechnol J. 2015 Jan 1;13:407–16.

7. Jiang X, Brantley Hall A, Arthur TD, Plichta DR, Covington CT, Poyet M, et al. Invertible promoters mediate bacterial phase variation, antibiotic resistance, and host adaptation in the gut. Science. 2019 Jan;363(6423):181–7.

8. Seib KL, Jen FEC, Scott AL, Tan A, Jennings MP. Phase variation of DNA methyltransferases and the regulation of virulence and immune evasion in the pathogenic Neisseria. Pathog Dis. 2017 Aug 31;75(6):ftx080.

9. Phillips ZN, Tram G, Seib KL, Atack JM. Phase-variable bacterial loci: how bacteria gamble to maximise fitness in changing environments. Biochem Soc Trans. 2019 Aug 30;47(4):1131–41.

10. Béchon N, Mihajlovic J, Vendrell-Fernández S, Chain F, Langella P, Beloin C, et al. Capsular Polysaccharide Cross-Regulation Modulates Bacteroides thetaiotaomicron Biofilm Formation. mBio. 2020 Jun 23;11(3):e00729–20.

11. Porter NT, Canales P, Peterson DA, Martens EC. A Subset of Polysaccharide Capsules in the Human Symbiont Bacteroides thetaiotaomicron Promote Increased Competitive Fitness in the Mouse Gut. Cell Host Microbe. 2017 Oct 11;22(4):494–506.e8.

12. Porter NT, Hryckowian AJ, Merrill BD, Fuentes JJ, Gardner JO, Glowacki RWP, et al. Phase-variable capsular polysaccharides and lipoproteins modify bacteriophage susceptibility in Bacteroides thetaiotaomicron. Nat Microbiol. 2020 Sep;5(9):1170–81.

13. Saunders NJ, Moxon ER, Gravenor MB. Mutation rates: estimating phase variation rates when fitness differences are present and their impact on population structure. Microbiology. 2003 Feb 1;149(2):485–95.

14. Criss AK, Kline KA, Seifert HS. The frequency and rate of pilin antigenic variation in Neisseria gonorrhoeae. Mol Microbiol. 2005 Oct;58(2):510–9.

15. Hung M, Chang E, Hussein R, Frazier K, Shin JE, Sagawa S, et al. Modulating the frequency and bias of stochastic switching to control phenotypic variation. Nat Commun. 2014 Aug 4;5(1):4574.

16. Krinos CM, Coyne MJ, Weinacht KG, Tzianabos AO, Kasper DL, Comstock LE. Extensive surface diversity of a commensal microorganism by multiple DNA inversions. Nature. 2001 Nov;414(6863):555–8.

17. Coyne MJ, Weinacht KG, Krinos CM, Comstock LE. Mpi recombinase globally modulates the surface architecture of a human commensal bacterium. Proc Natl Acad Sci. 2003 Sep 2;100(18):10446–51.

18. Johnson RC. Site-specific DNA Inversion by Serine Recombinases. Microbiol Spectr. 2015 Feb 19;3(3):1–36.

19. Lan F, Saba J, Ross TD, Zhou Z, Krauska K, Anantharaman K, et al. Massively parallel single-cell sequencing of genetic loci in diverse microbial populations [Internet]. bioRxiv; 2022 [cited 2022 Nov 28]. p. 2022.11.21.517444. Available from: https://www.biorxiv.org/content/10.1101/2022.11.21.517444v1

20. Kuwahara T, Yamashita A, Hirakawa H, Nakayama H, Toh H, Okada N, et al. Genomic analysis of Bacteroides fragilis reveals extensive DNA inversions regulating cell surface adaptation. Proc Natl Acad Sci. 2004 Oct 12;101(41):14919–24.

21. Qian Y, Lan F, Venturelli OS. Towards a deeper understanding of microbial communities: integrating experimental data with dynamic models. Curr Opin Microbiol. 2021 Aug 1;62:84–92.

22. Munsky B, Fox Z, Neuert G. Integrating Single-Molecule Experiments and Discrete Stochastic Models to Understand Heterogeneous Gene Transcription Dynamics. Methods San Diego Calif. 2015 Sep 1;85:12–21.

23. Liu CH, Lee SM, VanLare JM, Kasper DL, Mazmanian SK. Regulation of surface architecture by symbiotic bacteria mediates host colonization. Proc Natl Acad Sci. 2008 Mar 11;105(10):3951–6.

24. Coyne MJ, Tzianabos AO, Mallory BC, Carey VJ, Kasper DL, Comstock LE. Polysaccharide Biosynthesis Locus Required for Virulence of Bacteroides fragilis. Infect Immun. 2001 Jul;69(7):4342–50.

25. Troy EB, Carey VJ, Kasper DL, Comstock LE. Orientations of the Bacteroides fragilis Capsular Polysaccharide Biosynthesis Locus Promoters during Symbiosis and Infection. J Bacteriol. 2010 Nov;192(21):5832–6.

26. Patrick S, Parkhill J, McCoy LJ, Lennard N, Larkin MJ, Collins M, et al. Multiple inverted DNA repeats of Bacteroides fragilis that control polysaccharide antigenic variation are similar to the hin region inverted repeats of Salmonella typhimurium. Microbiology. 149(4):915–24.

27. Venturelli OS, Zuleta I, Murray RM, El-Samad H. Population Diversification in a Yeast Metabolic Program Promotes Anticipation of Environmental Shifts. PLOS Biol. 2015 Jan 27;13(1):e1002042.

28. Cowley SC, Myltseva SV, Nano FE. Phase variation in Francisella tularensis affecting intracellular growth, lipopolysaccharide antigenicity and nitric oxide production. Mol Microbiol. 1996 May;20(4):867–74.

29. Snellings NJ, Tall BD, Venkatesan MM. Characterization of Shigella type 1 fimbriae: expression, FimA sequence, and phase variation. Infect Immun. 1997 Jun;65(6):2462–7.

30. Lavitola A, Bucci C, Salvatore P, Maresca G, Bruni CB, Alifano P. Intracistronic transcription termination in polysialyltransferase gene (siaD) affects phase variation in Neisseria meningitidis. Mol Microbiol. 1999 Jul;33(1):119–27.

31. Snyder LA, Loman NJ, Linton JD, Langdon RR, Weinstock GM, Wren BW, et al. Simple sequence repeats in Helicobacter canadensis and their role in phase variable expression and C-terminal sequence switching. BMC Genomics. 2010;11(1):67.

32. Chopra-Dewasthaly R, Baumgartner M, Gamper E, Innerebner C, Zimmermann M, Schilcher F, et al. Role of Vpma phase variation in *Mycoplasma agalactiae* pathogenesis. FEMS Immunol Med Microbiol. 2012 Dec;66(3):307–22.

33. Beare PA, Jeffrey BM, Long CM, Martens CM, Heinzen RA. Genetic mechanisms of Coxiella burnetii lipopolysaccharide phase variation. Monack DM, editor. PLOS Pathog. 2018 Feb 26;14(3):e1006922.

34. Safi H, Gopal P, Lingaraju S, Ma S, Levine C, Dartois V, et al. Phase variation in *Mycobacterium tuberculosis glpK* produces transiently heritable drug tolerance. Proc Natl Acad Sci. 2019 Sep 24;116(39):19665–74.

35. Zhang AN, Li L, Yin X, Dai CL, Groussin M, Poyet M, et al. Choosing Your Battles: Which Resistance Genes Warrant Global Action? bioRxiv. 2019 Oct;(October):784322.

36. Duffy DC, McDonald JC, Schueller OJA, Whitesides GM. Rapid Prototyping of Microfluidic Systems in Poly(dimethylsiloxane). Anal Chem. 1998 Dec;70(23):4974–84.

37. Langmead B, Salzberg SL. Fast gapped-read alignment with Bowtie 2. Nat Methods. 2012 Apr;9(4):357–9.

