## Supplementary Information for "High-throughput single-cell sequencing of multiple invertible promoters reveals a strong determinant of bacterial population heterogeneity"

#### A mathematical model of stochastic phase variation

Here we develop a mathematical model to describe the dynamic evolution of CPS promoter heterogeneity in a population of *B. fragilis*. Such heterogeneity at the population level is a consequence of the inherent stochastic nature of biomolecular reactions occurring at the single cell level. In particular, we assume that the dynamic processes driving CPS promoter inversion have the following ergodicity property: at any time $t$ and for any promoter state $i$, the *fraction* of cells in state $i$ in a population is equal to the *probability* of a single cell to be in state $i$. Hence, in the sequel, we develop a mathematical model to describe the stochastic biomolecular reactions at the single cell level that governs CPS promoter inversion.

In each *B. fragilis* cell, there are $n = 7$ CPS gene clusters. Let $\mathrm{x}_i$ represent the $i$-th promoter, which is either in "ON" state (denoted by $\mathrm{x}_i = 1$) or in "OFF" state (denoted by $\mathrm{x}_i = 0$). When an invertase $E$ binds with the promoter $i$, it could switch the promoter to the "ON" ("OFF") state if it was previously in the "OFF" ("ON") state. We use the following chemical reaction to describe the binding and unbinding between the invertase and the promoter $i$, as well as flipping of the promoter state:

$$(\mathrm{x_i} = 0) + \mathrm{E} \rightleftharpoons (\mathrm{x_i} = 1) + \mathrm{E}, \quad \text{for every promoter } i. \tag{1}$$

The *macrostate* of a cell is dictated by the combination of its CPS promoter states and is denoted by a *random variable* z. With $n$ invertible CPS promoters, this constitutes a $2^n$ dimensional discrete state space $\mathrm{z} \in \mathcal{Z} = \{0, 1\}^n$. To denote possible *realizations* of system macrostates, we will use $z_i$, where $i$ is the decimal conversion of the binary sequence representing promoter states. For example, $\mathrm{z} = z_0 := (0000000)$ denotes the cell is in a macrostate where all promoters are "OFF" and $\mathrm{z} = z_{127} := (1111111)$ denotes the cell is in a macrostate where all promoters are "ON". For a macrostate $z_i$, a different macrostate $z_j$ is its *neighbor state* if they are different by a *single* promoter inversion. We use $\mathcal{N}_i$ to denote the set of all neighbor states of $z_i$. For instance, the set of neighbors of state $z_0$ is $\mathcal{N}_0 = \{z_1, z_2, z_4, z_8, z_{16}, z_{32}, z_{64}\}$. For every $i$, we have $|\mathcal{N}_i| = n = 7$. Now we develop a mathematical model to describe the stochastic transition between cell macrostates. The resultant model is a set of coupled linear ordinary differential equations (ODEs) describing how the probability of the cell in each macrostate $i$, denoted as $P_i(t) := P(\mathrm{z} = z_i, t)$, evolves over time. Such a model is known as a *chemical master equation* model in the chemical reaction network setting [1, 2], and here we derive it in full detail to explicitly account for all relevant physical assumptions.

We first consider generic state transition dynamics in a continuous-time discrete-state Markovian setting. Taking a sufficiently small time interval $\delta t$, in which only a *single* state transition (i.e., chemical reaction) could occur. The probability a cell is in state $z_i \in \mathcal{Z}$ at time $t + \delta t$ is determined by the propensities of the following two types of events:

(I) The propensity $\alpha_{i,j}\delta t$ that the cell starts at a neighbor state $z_j \in \mathcal{N}_i$ at time $t$, but transition to $z_i$ by inverting a single promoter during the $\delta t$ interval.

(II) The propensity that the cell starts at $z_i$ at $t$ and no reaction occur during the $\delta t$ interval. This propensity can be quantified as $(1 - \sum_{j \in \mathcal{N}_i} \alpha_{j,i}\delta t)$, where $\alpha_{j,i}$ is the propensity of the cell in state $i$ to transition to a neighbor state $j \in \mathcal{N}_i$.

We can then compute the probability of each cell to be in state $z_i$ at time $t + \delta t$ as follows:

$$P_i(t + \delta t) = P_i(t)[1 - \sum_{j \in \mathcal{N}_i} \alpha_{j,i}\delta t] + \sum_{j \in \mathcal{N}_i} P_j(t)\alpha_{i,j}\delta t, \quad \text{for all } \; i = 1, \cdots, 127. \tag{2}$$

Rearranging equation (2), we have

$$\frac{\mathrm{d}}{\mathrm{d}t} P_i(t) := \lim_{\delta t \to 0^+} \frac{P_i(t + \delta t) - P_i(t)}{\delta t} = \sum_{j \in \mathcal{N}_i} \alpha_{i,j} P_j(t) - \sum_{j \in \mathcal{N}_i} \alpha_{j,i} P_i(t), \quad \text{for all } \; i = 1, \cdots, 127. \tag{3}$$

Alternatively, equation (3) can be put in matrix form:

$$\frac{\mathrm{d}}{\mathrm{d}t}\begin{bmatrix} P_0 \\ P_1 \\ \vdots \\ P_{127} \end{bmatrix} = \underbrace{\begin{bmatrix} -\sum_{j\in\mathcal{N}_0}\alpha_{j,0} & \alpha_{0,1} & \cdots & \alpha_{0,127} \\ \alpha_{1,0} & -\sum_{j\in\mathcal{N}_1}\alpha_{j,1} & \cdots & \alpha_{1,127} \\ \vdots & \vdots & \ddots & \vdots \\ \alpha_{127,0} & \alpha_{127,1} & \cdots & -\sum_{j\in\mathcal{N}_{127}}\alpha_{j,127} \end{bmatrix}}_{A} \begin{bmatrix} P_0 \\ P_1 \\ \vdots \\ P_{127} \end{bmatrix}. \tag{4}$$

Since equation (3) is linear, its solution can be written as

$$\mathbf{P}(t) = \exp(A\cdot t)\cdot \mathbf{P}(0), \tag{5}$$

where $\mathbf{P} := [P_1, \cdots, P_{127}]^\top$, $A$ is called the *state transition matrix*, and $\exp(\cdot)$ represents matrix exponential operation.

We then write the parameters in the transition matrix $A$ in terms of parameters in the chemical reaction (1). To this end, we consider the probability that each reaction in (1) occurs. In particular, for each promoter $i$ and each forward (or backward) reaction in (1), the *reaction propensity functions* $a_i^f(\mathrm{z}, t)\delta t$ (or $a_i^b(\mathrm{z}, t)\delta t$, respectively) captures the instantaneous probability that the reaction occurs (i.e., the promoter flips). More specifically, the reaction propensity function is the probability the forward (or backward) reaction will occur for promoter $i$ between $t$ and $t+\delta t$ given current state z and an infinitesimal time $\delta t$ [2]. Assuming that the cell is a well-mixed space, the probability that an invertase collides and reacts with promoter $i$ in "ON" (or "OFF") state is determined by the following factors [1, 2]: (i) it is proportional to the number of invertase molecules (ii) proportional to the number (*i.e.*, 0 or 1) of promoter $i$ in "ON" (or "OFF") state, and (iii) it is inversely proportional to the volume of the cell $\Omega$. Hence, the reaction propensity functions for the forward and backward reactions can be written as:

$$a_i^f(\mathrm{z}) = \frac{1}{\Omega}\cdot r_i^+ \cdot n_E \cdot \mathbb{1}(\mathrm{x}_i = 0), \qquad a_i^b(\mathrm{z}) = \frac{1}{\Omega}\cdot r_i^- \cdot n_E \cdot \mathbb{1}(\mathrm{x}_i = 1), \tag{6}$$

where $n_E$ is the number of invertase molecules, $r_i^+/r_i^-$ are the forward/backward reaction rate constants, and $\mathbb{1}(u)$ is the indicator function with $\mathbb{1}(u) = 1$ if $u$ is true and $\mathbb{1}(u) = 0$ if otherwise.

We consider transition from state $z_j$ to $z_i$, quantified by the propensity function $\alpha_{i,j}\delta t$. Depending on the promoter configurations in states $z_j$ and $z_i$, there are three cases, and every entry in the transition matrix $A$ can be determined by the following rules:

(a) If $i \notin \mathcal{N}_j$, based on equation (3), we have:

$$\alpha_{i,j} = 0. \tag{7}$$

(b) If $i \in \mathcal{N}_j$ and the $q$-th promoter ($q = 1, \cdots, 7$) is switched from "OFF" in $z_j$ to "ON" in $z_i$ in $\delta t$ time interval, then according to (6), we have:

$$\alpha_{i,j} = a_q^f(z_j) = r_q^+ \cdot [E], \tag{8}$$

where $[E]$ denotes the concentration of the invertase E.

(c) Conversely, if $i \in \mathcal{N}_j$ and the $q$-th promoter is switched from "ON" in state $z_j$ to "OFF" in state $z_i$ in $\delta t$ time interval, then according to (6), we have:

$$\alpha_{i,j} = a_q^b(z_j) = r_q^- \cdot [E], \tag{9}$$

where $[E]$ denotes the concentration of the invertase E.

Here, we further assume that the invertase is expressed in sufficiently large amount such that its copy number is much larger than 7 (i.e., the total number of promoter sites it could bind to). As a result, the free invertase concentration $[E]$ stays approximately constant with individual promoter inversion events. Under this assumption, we call $\hat{r}_i^\pm := r_i^\pm \cdot [E]$ the *effective flipping rates*. Hence, in sum, the propensity of a cell switching from a macrostate $z_j$ to a neighbor state $z_i$ (i.e., $\alpha_{i,j}$) by flipping a promoter is equal to the corresponding effective flipping rate, which we determine from experimental data in the next section.

#### Parameter inference

To infer the effective flipping rates $\hat{r}_i^{\pm}$ as well as their associated uncertainties from experimental data, we perform Markov Chain Monte Carlo (MCMC) simulations. This allows us to obtain posterior distributions of the parameters $\theta := [\hat{r}_1^+, \hat{r}_1^-, \cdots, \hat{r}_7^+, \hat{r}_7^-]$ given experimentally measured *empirical distribution* dynamics starting from different initial conditions. Specifically, let $N_i(t,k)$ be the number of cells in macrostate $z_i$ measured at time $t$ with initial condition $k$, the respective empirical distribution $\hat{\mathbf{P}}(t,k)$ is defined as:

$$\hat{P}_i(t,k) := \frac{N_i(t,k)}{\sum_{j=0}^{127} N_j(t,k)}, \tag{10}$$

with $\hat{\mathbf{P}}(t,k) := [\hat{P}_0(t,k), \cdots, \hat{P}_{127}(t,k)]$. We intend to find parameters that allow $\mathbf{P}$ dynamics in (5) to match the evolution of empirical distributions $\hat{\mathbf{P}}$ starting from all initial conditions. We use an additive, Gaussian noise to model measurement uncertainty:

$$\hat{P}_i(t,k) = P_i(t,k) + \varepsilon_i, \text{ where } \varepsilon_i \sim \mathcal{N}(0, \sigma_i(t,k)) \tag{11}$$

and $\sigma_i$ is the standard deviation associated with the fraction of cells in state $z_i$. For states with smaller number of cell counts, the deviation between probability $P_i$ and empirical distribution $\hat{P}_i$ increases. To account for this, we further assume that measurement uncertainty increases for states with low cell counts and hence fractions: $\sigma_i(t,k) = -a \cdot \log_{10} \hat{P}_i(t,k) + b$. The parameters $a = 0.028$ and $b = 0.016$ are found by fitting standard deviation quantified from three independent DoTA-seq experiments. Given mechanistic state transition model (5) and noise model (11), we use $\hat{\mathbf{X}} := [\hat{\mathbf{P}}(t_1,k_1), \cdots, \hat{\mathbf{P}}(t_m,k_1), \hat{\mathbf{P}}(t_1,k_2), \cdots, \hat{\mathbf{P}}(t_m,k_q)]$ to represent the measured time series of cell fractions for all macrostates starting from all initial conditions $k_1, \cdots, k_q$. For a fixed parameter $\theta$, the likelihood to observe $\hat{\mathbf{X}}$ can be computed as

$$\mathbb{P}(\hat{\mathbf{X}}|\theta) = \prod_{j=1}^{m} \prod_{l=1}^{q} \prod_{i=0}^{127} f(\hat{P}_i(t_j,k_l) - P_i(t_j,k_l); \sigma_i(t_j,k_l)), \tag{12}$$

where $f(\cdot;\sigma)$ is the probability density function for the normal distribution with standard deviation $\sigma$ and $P_i(t_j,k_l)$ is the solution to (5) from the $l$-th initial condition (i.e., setting the empirical distribution measured at $t = 0$ from the $l$-th initial condition as $\mathbf{P}(0)$). The posterior probability can then be described according to Bayes rule as $\mathbb{P}(\theta|\hat{\mathbf{X}}) \propto \mathbb{P}(\hat{\mathbf{X}}|\theta) \cdot \pi(\theta)$, where $\pi(\theta)$ is the prior parameter distribution. We assign a uniform prior distribution $U(0,10)$ for each parameter.

An adaptive, symmetric, random-walk Metropolis MCMC algorithm [3] is then used to draw samples from the posterior distributions. Implementation details of the same MCMC setup has been described in [4, 5]. The algorithm was implemented using custom code in MATLAB R2021a (The MathWorks, Inc., Natick, MA, USA). For each parameter, we collected at least 100,000 MCMC samples after a burn-in. The Gelman-Rubin potential scale reduction factor (PSRF) was used to evalute convergence of the posterior distribution, where a PSRF closer to 1 indicates better convergence. The PSRF for all parameters lie between 1.0002 and 1.0056. In order to verify that the model structure in (5) is not too flexible to fit any experimental data, we randomly shuffled experimental data by exchanging the measured fractions of cells in each state. In particular, we generate randomly shuffled data $\tilde{\mathbf{P}}(t,k)$ via $\tilde{\mathbf{P}}(t,k) = \Lambda \cdot \hat{\mathbf{P}}(t,k)$, where $\Lambda \in \mathbb{R}^{128\times128}$ randomly shuffles elements in $\hat{\mathbf{P}}(t,k)$ (i.e., it contains a one at a random position in each row and each column only contains one non-zero element) and it is constant for all initial condition and all time. For both the synthetic strain data and WT data, we generated 100 such $\Lambda$ matrices hence 100 sets of randomly shuffled data. We use Pearson correlation between $\mathbf{P}(t,k)$ and $\hat{\mathbf{P}}(t,k)$ for all $t$ and $k$ to evaluate goodness of fit and found that the mechanistic model (5) had difficulty fitting randomly shuffled data. Specifically, for the synthetic strain (WT) data, Pearson correlation for all 100 sets of randomly shuffled data are below 0.6 (0.61, for WT respectively). In comparison, Pearson correlations for the unshuffled data were 0.97 and 0.80 for the synthetic strain and the WT data, respectively. These results indicate that our model (5) is not too flexible to fit randomly generate data hence it partially explains the experimental observations.

### References


[1] N. G. Van Kampen. *Stochastic Processes in Physics and Chemistry.* Elsevier, 2007.

[2] D. Del Vecchio and R. M. Murray. *Biomolecular Feedback Systems.* Princeton University Press, Princeton, 2014.

[3] Heikki Haario, Eero Saksman, and Johanna Tamminen. An adaptive metropolis algorithm. *Bernoulli*, 7(2):223, 2001.

[4] Susan Hromada, Yili Qian, Tyler B Jacobson, Ryan L Clark, Lauren Watson, Nasia Safdar, Daniel Amador-Noguez, and Ophelia S Venturelli. Negative interactions determine *Clostridioides difficile* growth in synthetic human gut communities. *Molecular Systems Biology*, 17(10), 2021.

[5] Jun Feng, Yili Qian, Zhichao Zhou, Sarah Ertmer, Eugenio I. Vivas, Freeman Lan, Joshua J. Hamilton, Federico E. Rey, Karthik Anantharaman, and Ophelia S. Venturelli. Polysaccharide utilization loci in bacteroides determine population fitness and community-level interactions. *Cell Host and Microbe*, 30(2):200–215.e12, 2022.

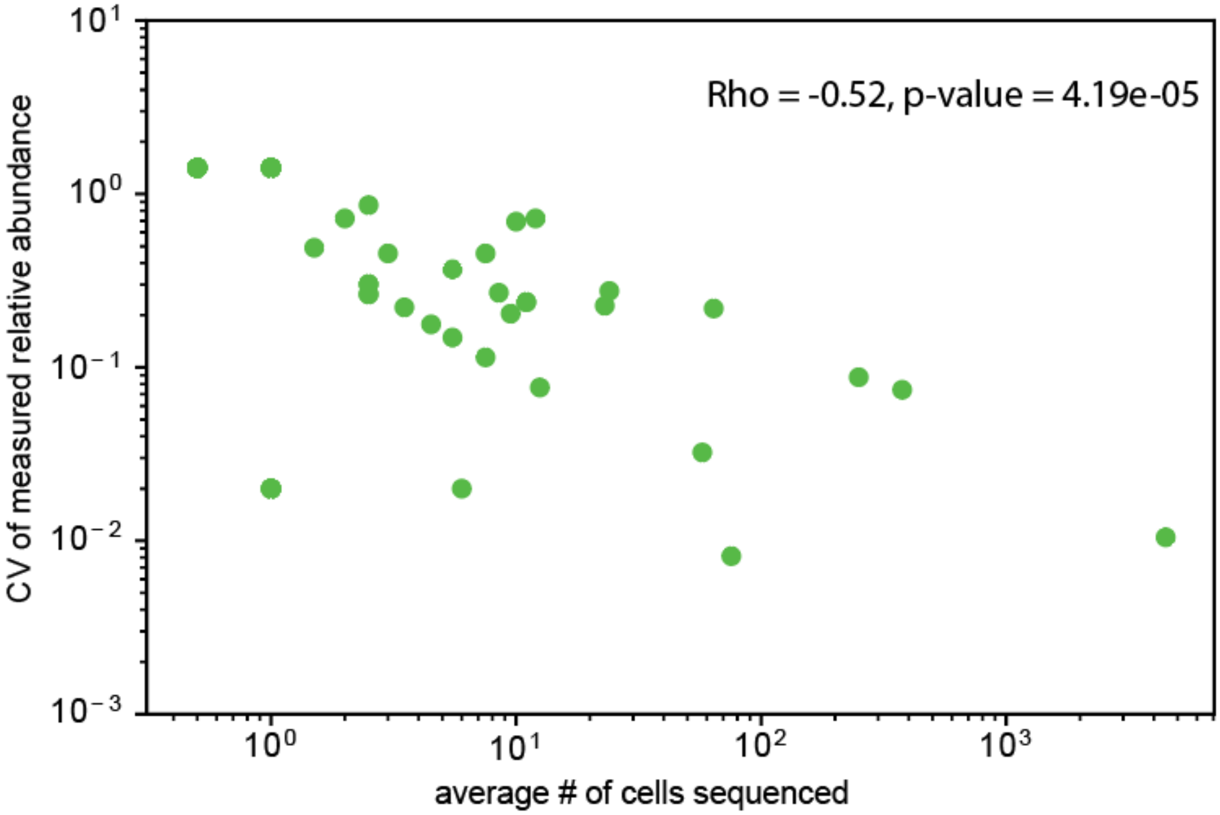


**Supplementary Figure 1**. Coefficient of variation (CV) of the measured relative abundance is negatively correlated with the number of cells sequenced for each subpopulation. Rho represents the Spearman's correlation coefficient for the dataset.

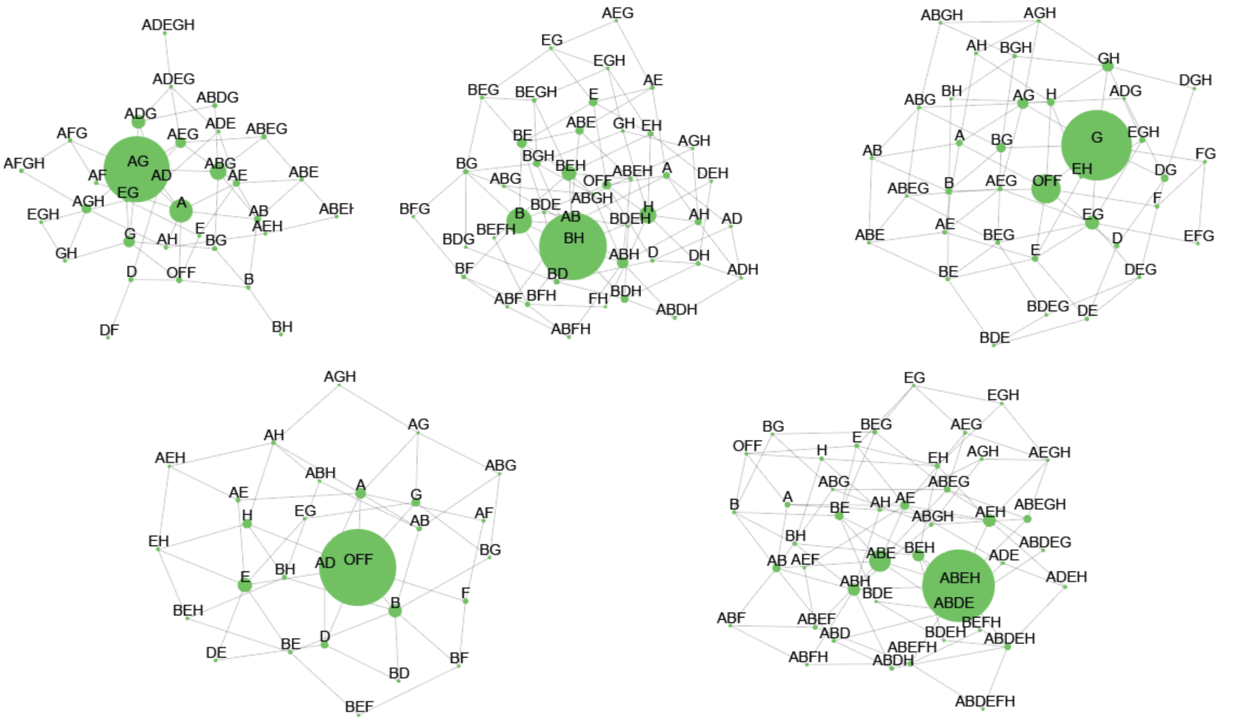


**Supplementary Figure 2.** Network graphs representing the population structures of different colonies grown, as determined by DoTA-seq. Each node represents a different subpopulation with the label indicating the combination of promoters turned ON. The "OFF" label represents subpopulations with all invertible promoters turned OFF. Node sizes represent the relative population abundances and edges connect nodes that are one promoter flip away.

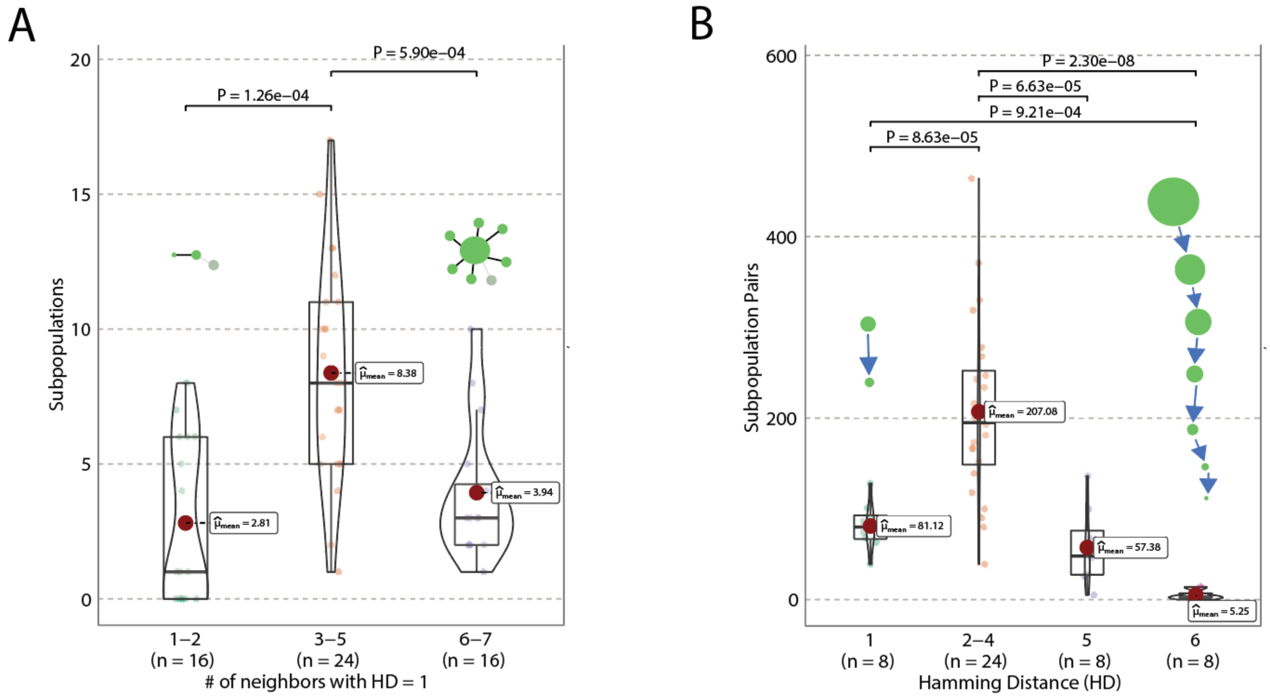


**Supplementary Figure 3.** Summary statistics for all colonies.

A) Frequency distributions as violin plot of immediate (Hamming Distance (HD) = 1) neighbor count for all subpopulations within their respective colonies. A boxplot is overlayed and the mean highlighted as a red dot. Welch's ANOVA was performed and supplemented by post-hoc pairwise Games-Howell tests. Holm-Bonferroni-corrected p-values are reported for pairwise comparisons. B) Frequency distribution as violin plot of HDs for all possible pairs of subpopulations within their respective colonies. A boxplot is overlayed and the mean highlighted as a red dot. Welch's ANOVA was performed and supplemented by post-hoc pairwise Games-Howell tests. Holm-Bonferroni-corrected p-values are reported for pairwise comparisons. Horizontal brackets represent statistical significant pairwise comparisons.

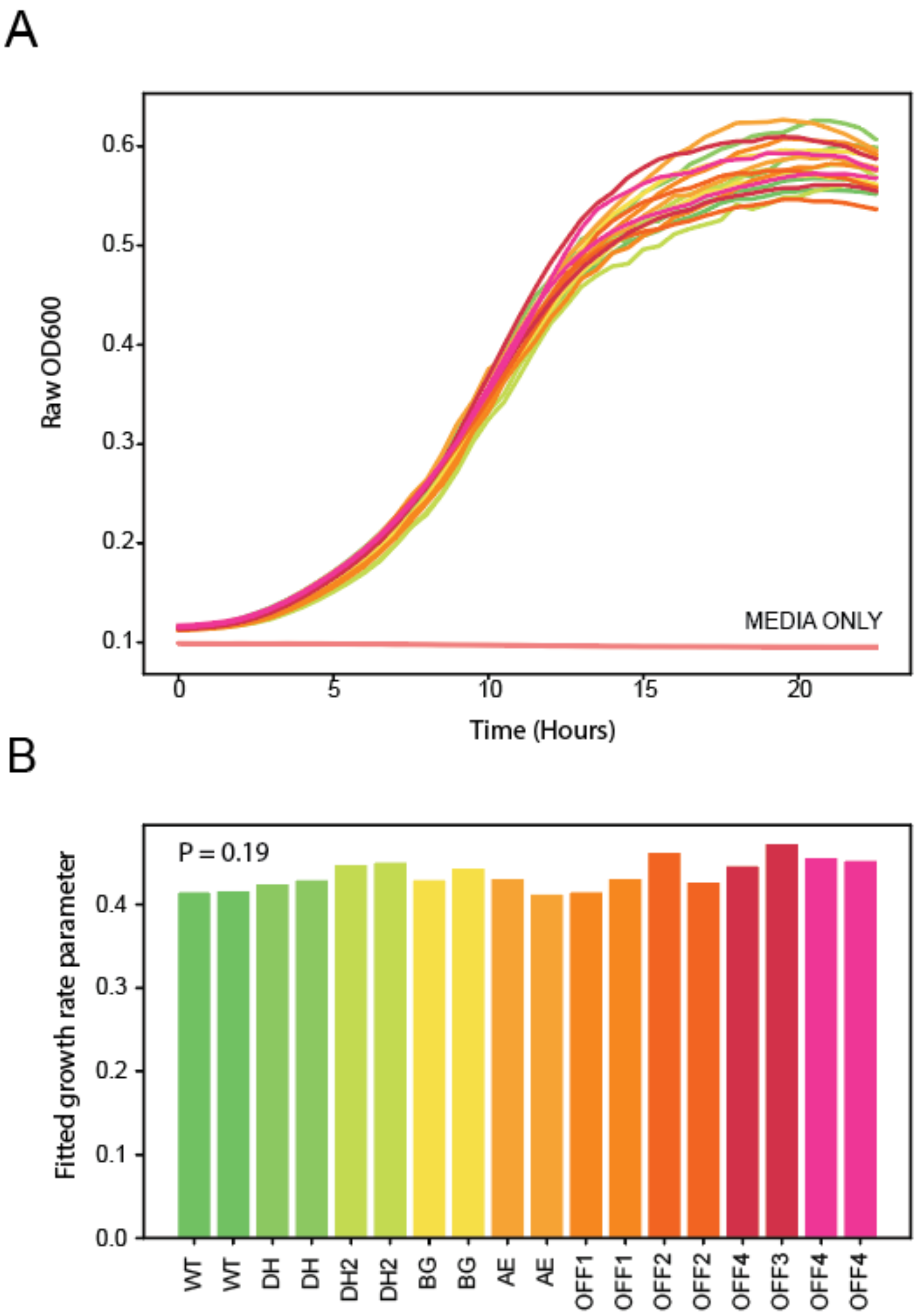


**Supplementary Figure 4.** Growth rates are similar between different CPS locked strains. A) Raw OD600 growth curves of wildtype and different CPS locked strains grown in media. B) Growth rate parameters of wildtype (WT) and strains locked in different CPS promoter orientations as determined by growth curve fitting to a logistic growth model. Each strain (unique label on the X axis) is grown in technical duplicates. One-factor ANOVA between the different promoter orientations shows no statistically significant difference (P-value 0.19) between the growth rates among the different CPS promoter orientations. Colors indicate different strains tested and are consistent across A and B.

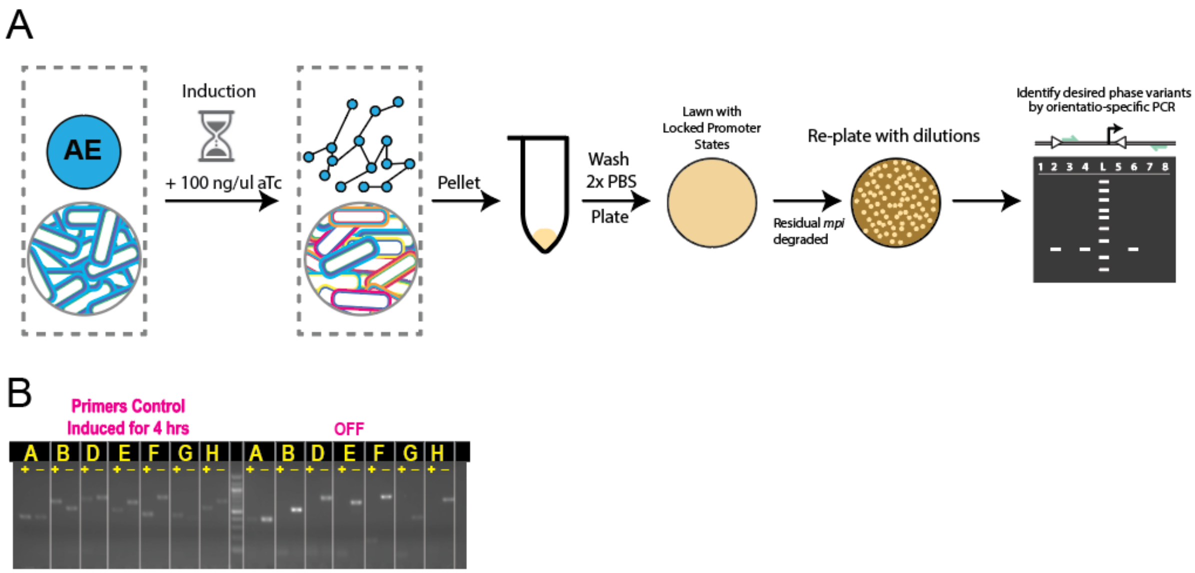


**Supplementary Figure 5.** Workflow for rapidly isolating and characterizing phase-locked variants. a) Schematic of workflow for generating different phase-locked variants. Starting with an inducible invertase strain, after 4 hours of induction, a population with diverse combinations of promoter orientations was generated. Cells were washed twice to remove residual amounts of inducer, then plated for overnight growth and gradual dilution and degradation of residual invertase. The population is re-streaked to new plates to isolate single colonies largely fixed in promoter orientations. b) Representative results of orientation-specific PCR using previously published primers (Coyne et al., 2003), which is used to identify the promoter orientations of different locked colonies. The '+' signifies a lane in which PCR was done using primers to detect an on-oriented promoter, while conversely the '-' indicates a lane in which PCR was done using primers to detect an off-oriented promoter. The extreme sensitivity of orientation-specific PCR suggests a small fraction of PSA on, and this sensitivity was noted previously with these published primer sets (Coyne et al 2003).

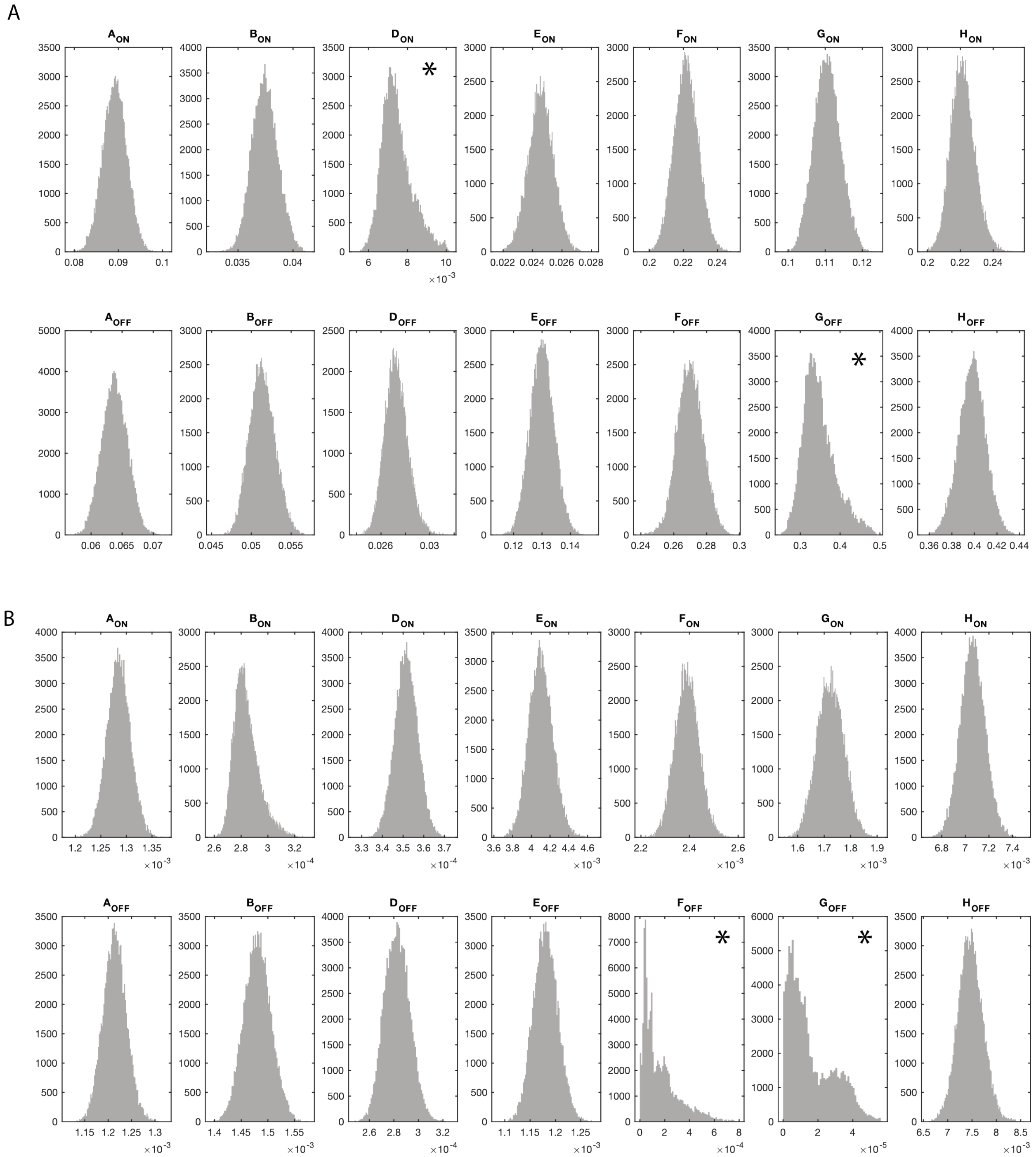


**Supplementary Figure 6**. Parameter distributions obtained from Markov Chain Monte Carlo fitting of the stochastic inversion model on induced population timeseries data (A) and wildtype population timeseries data (B). Asterix (*) indicates parameters where the coefficient of variation is larger than 5%, indicating possible insufficient constraint on those parameters.

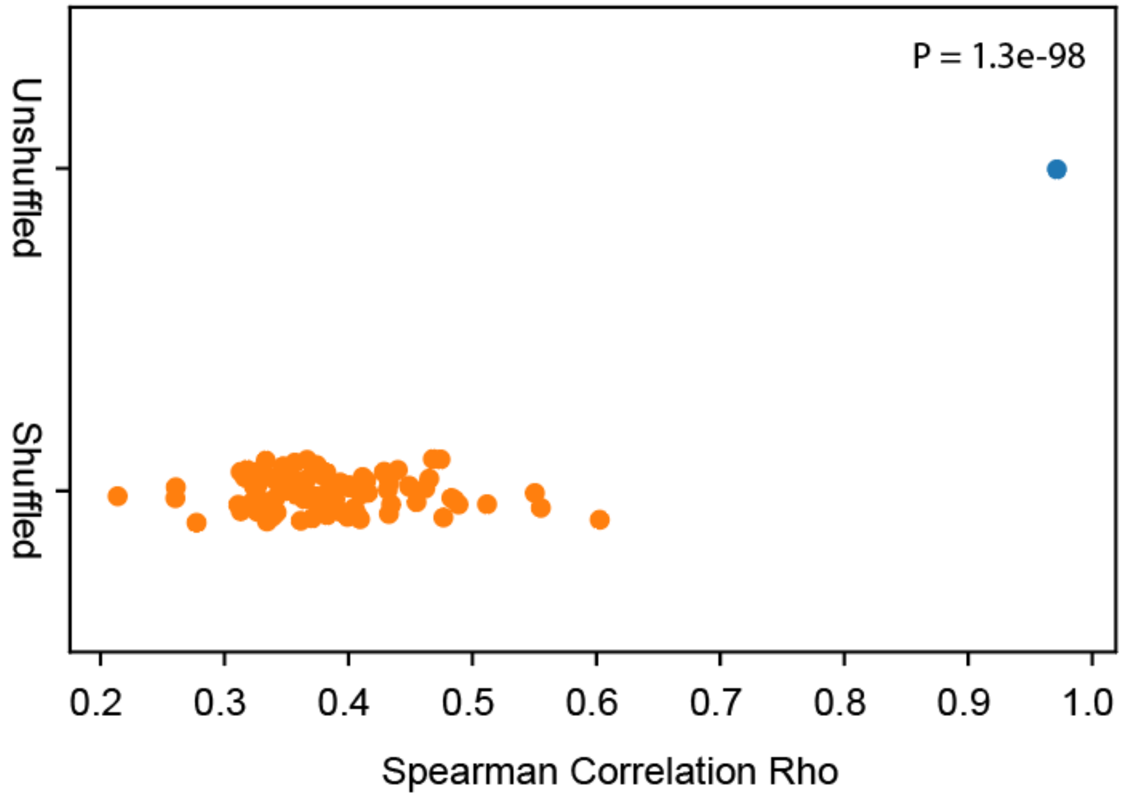


**Supplementary Figure 7.** Spearman correlation between experimental data and model predictions for models generated using unshuffled and label shuffled experimental data from the induced invertase timeseries. Experimental data with unshuffled and 100 iterations of randomly shuffled labels are fit to the promoter inversion model using Markov-Chain Monte Carlo (see Methods). Spearman's correlation between the model predictions and the data used to fit the models is used as a measure of goodness-of-fit. Unshuffled data obtains significantly higher correlation (Rho = 0.98) compared to shuffled data controls (Rho = 0.38 +/- 0.06) showing that the model data agreement would not arise from random data.

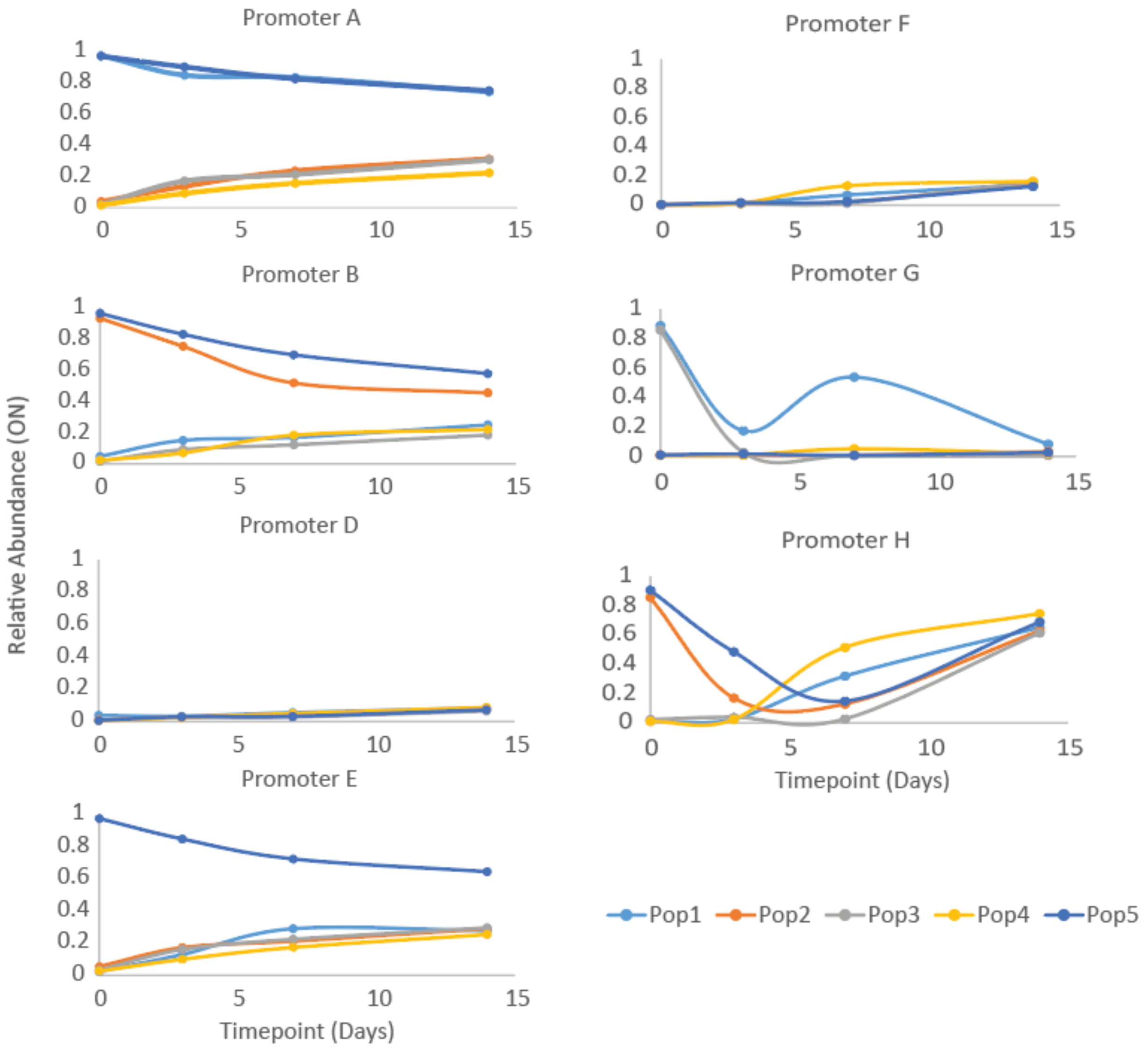


**Supplementary Figure 8.** Diversification trajectories of each promoter within the 5 wild-type populations. Promoters A-E appear to converge at a slower rate than promoters F-H.
